# PD-1 Specific ‘Blocking’ Antibodies That Deplete PD-1+ T Cells Present An Inconvenient Variable In Pre-clinical Immunotherapy Experiments

**DOI:** 10.1101/2020.04.14.041608

**Authors:** Fanny Polesso, Michael W. Munks, Katherine H. Rott, Savannah Smart, Ann B. Hill, Amy E. Moran

## Abstract

Therapeutic antibodies blocking PD-1-/PD-L1 interaction have achieved remarkable clinical success in cancer. In addition to blocking a target molecule, some isotypes of antibodies can activate complement, NK cells or phagocytes, resulting in death of the cell expressing the antibody’s target. Human anti-PD-1 therapeutics use antibody isotypes designed to minimize such antibody-dependent lysis. In contrast, anti-PD-1 reagents used in mice are derived from multiple species, with different isotypes, and are not engineered to reduce target cell death: few studies analyze or discuss how antibody species and isotype may impact data interpretation. We demonstrate here that anti-PD-1 therapy to promote activation and proliferation of PD-1-expressing CD8 T cells sometimes led instead to a loss of antigen specific cells. This phenomenon was seen in two tumor models and a model of virus infection, and was varied with the clone of anti-PD-1 antibody. Additionally, we compared competition among anti-PD-1 clones to find a combination that allows detection of PD-1-expressing cells despite the presence of blocking anti-PD1 antibodies *in vivo.* These data bring attention to the possibility of unintended target cell depletion with some commonly used anti-mouse PD-1 clones, and should provide a valuable resource for the design and interpretation of anti-PD-1 studies in mice. (200)

## Introduction

PD-1 is transiently expressed on T cells following T cell receptor activation. In some contexts of chronic antigen stimulation, for example within tumors or with some persistent viral infections, T cells become dysfunctional and PD-1 expression remains high. Inhibition of PD-1 signaling, through blockade with anti-PD-1 or anti-PD-L1, can improve T cell function (Barber et al., 2006; Pardoll, 2012), and has demonstrated widespread clinical success in cancer immunotherapy. Many clinical studies are now examining how best to combine PD-1 inhibition with traditional cancer treatment strategies, such as surgery, radiation, chemotherapy and targeted therapies.

To predict which combinations will provide synergy, there is immense interest in developing and characterizing genetically engineered mouse (GEM) models that mimic the most salient features of particular human cancers. In contrast, very little attention has been given to how research-grade anti-PD-1 reagents compare to human PD-1 therapeutics. For instance, nivolumab is a fully human anti-PD-1 IgG4, while pembrolizumab is a humanized anti-PD-1 IgG4. Human IgG4 lacks high-affinity FcγR binding and has low complement activity (Jefferis, 2012), thus nivolumab and pembrolizumab are expected to have minimal antibody-dependent cellular cytotoxicity (ADCC), antibody-dependent cellular phagocytosis (ADCP), and complement-dependent cytotoxicity (CDC). For nivolumab, a lack of effector activity has been directly demonstrated (Wang et al., 2014), and the lack of effector function in these IgG4 isotypes is critical for reducing or preventing the deletion of PD-1 positive T cells that could be tumor-antigen reactive. In contrast, mouse PD-1 antibodies are primarily either rat IgG2 (e.g. 29F.1A12, RMP1-14, RMP1-30) or hamster IgG (e.g. J43, G4), and we are unaware of studies characterizing whether or not they have effector functions mediated through either FcγR or complement.

While conducting immunotherapy experiments in mice, we observed profound differences between anti-PD-1 and anti-PD-L1. While anti-PD-L1 synergized with OX40 agonism (Polesso et al., 2019), anti-PD-1 attenuated OX40 immunotherapy (Shrimali et al., 2017). We show here that anti-PD-1 clone G4 (Armenian hamster IgG), when compared to anti-PD-1 clone RMP1-14 (rat IgG2a), or anti-PD-L1 clone 10F.9G2 (rat IgG2b), resulted in a loss of antigen-specific CD8 T cells in two sarcoma tumor models and the murine cytomegalovirus (MCMV) infection model. Our data indicate that anti-PD-1 can cause CD8 T cell depletion in vivo. To improve interpretation of experiments using anti-mouse PD-1, we propose more rigorous characterization of these reagents, and propose that rat and hamster PD-1 reagents could be expressed on a mouse IgG backbone lacking effector function.

## Results

### Anti-tumor immunity is enhanced with anti-PD-L1 but not anti-PD-1

In the MCA-205 mouse sarcoma model, we previously published that anti-PD-L1 synergizes with OX40 agonism to enhance anti-tumor T cell responses and cause tumor regression (Polesso et al., 2019). While performing these experiments, we compared anti-PD-1 to anti-PD-L1 immunotherapy (Fig 1A) and noted a striking difference. Anti-PD-1 led to animal death with nearly identical kinetics as the control treated animals (Fig 1 B-D). In contrast, 60% of anti-PD-L1 treated animals rejected their tumors (Fig 1B and E). The lack of therapeutic efficacy with anti-PD-1 is consistent with published data using the same clone of PD-1 antibody (clone G4) (Hirano et al., 2005), in which no therapeutic benefit was noted when treating tumor-bearing animals with this single agent (Messenheimer et al., 2017). We next treated mice with large tumors (~50-70 mm^2^) with a combination of agonist anti-OX40 and either anti-PD-1 or anti-PD-L1. Single-agent therapy did not provide a survival advantage (Supplemental Fig 1A and B). However, anti-PD-L1 synergized with anti-OX40 and led to tumor protection in 50% of animals, whereas anti-PD-1 did not synergize with anti-OX40, with no survival benefit compared to the control group (Supplemental Fig 1B-G). These results highlight a fundamental difference between the use of anti-PD-1 versus anti-PD-L1 in the success of single or combination immunotherapy in our MCA-205 tumor model.

**Figure 1.**
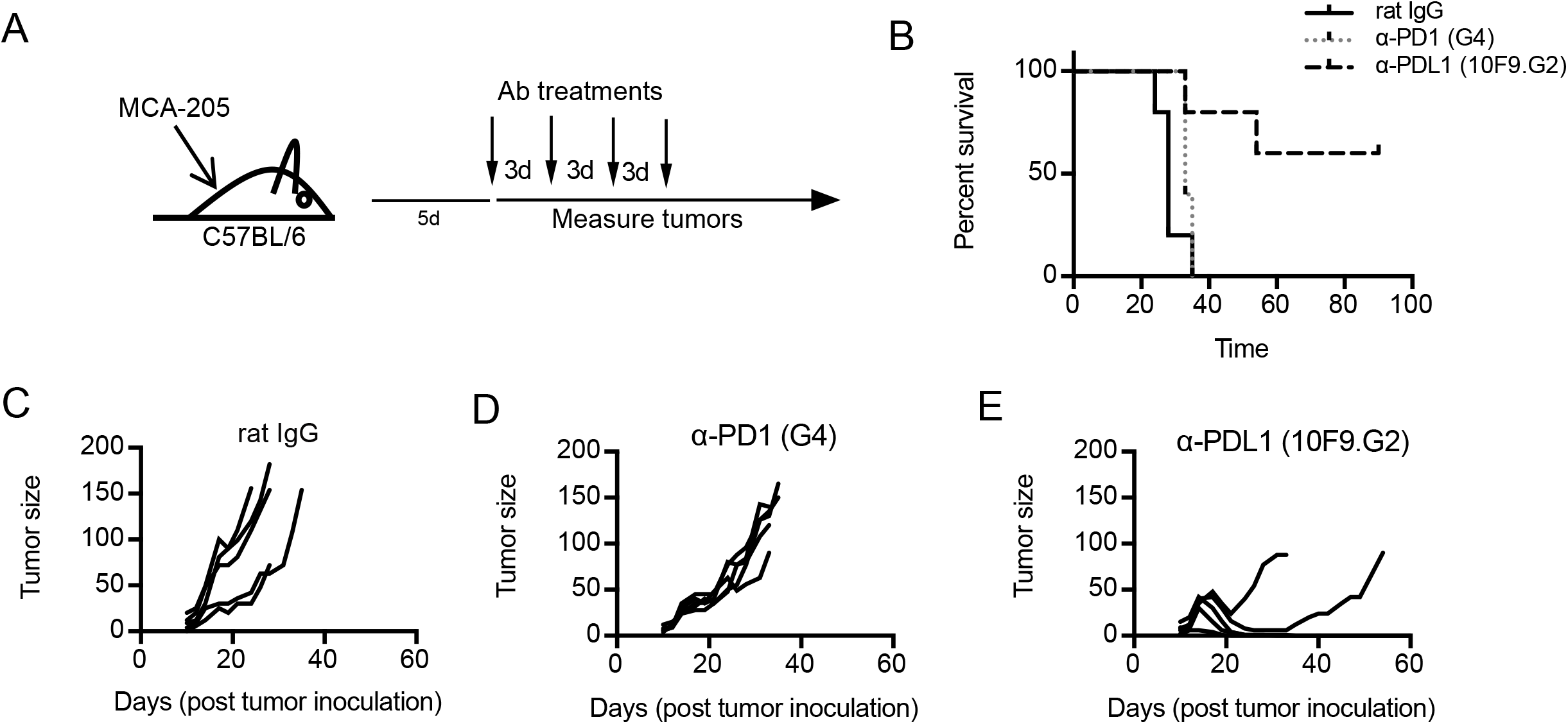
Survival outcomes are different in anti-PD-1 vs anti-PD-L1 treated animals. A) Experimental design of tumor experiments. Animals were treated with 200ug of rat IgG control, α-PD-1 or α-PD-L1. Tumors were measured 3 times a week and animals were euthanized when the tumors were >150mm2 or when tumors became ulcerated. B) Overall survival of MCA205 tumor-bearing C57Bl/6 mice. C-E) Individual tumor growth curves for each treatment group. Data are representative of more than 3 independent experiments with 5-7 animals per treatment group.

### PD-1 but not PD-L1 blockade leads to a decrease in tumor-antigen specific CD8 T cells

Given the diminished anti-tumor effects observed with anti-PD-1 compared to anti-PD-L1 combination therapy, we wondered if anti-PD-1 antibody reduced tumor-antigen specific T cell responses. Therefore, using a tetramer for mAlg8, a known tumor rejection antigen (Gubin et al., 2014), we analyzed tumor-antigen specific T cells from d42m1-T3 MCA sarcoma tumor-bearing animals and evaluated the expression of PD-1 (Fig 2A) and PD-L1 (Fig 2B) on T cells in the tumor and spleen. As previously reported (Polesso et al., 2019), tumor-antigen specific T cells in the tumor expressed very high levels of PD-1 compared to their counterpart in the spleen, and this increase was much greater than the expression of PD-L1 on these same cells.

**Figure 2.**
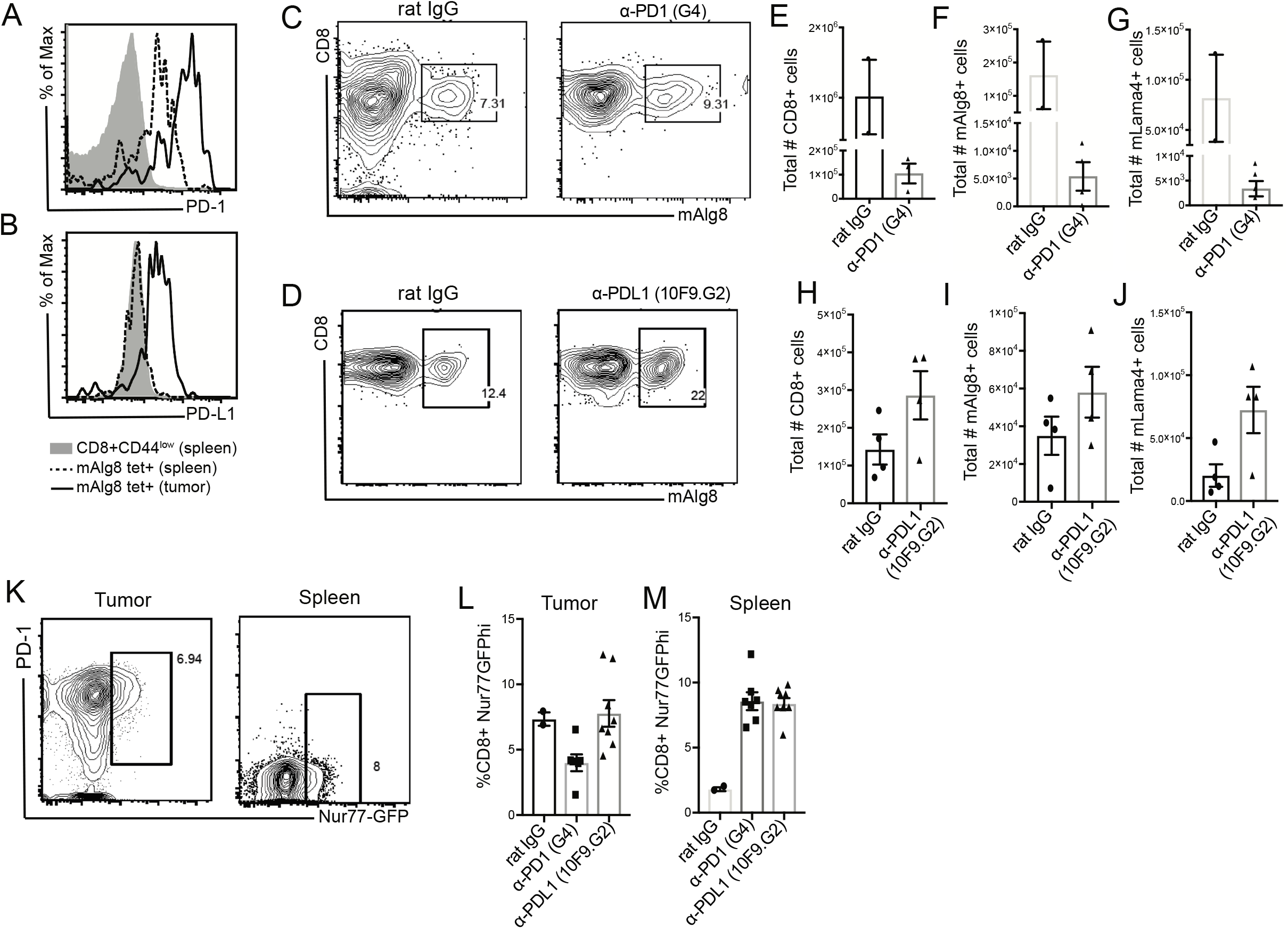
Tumor antigen specific T cells are diminished after treatment with anti-PD-1 but not anti-PD-L1. A) PD-1 and B) PD-L1 expression on tumor antigen specific, mAlg8 tetramer positive T cells in d42m1 tumors and spleen d17 after tumor inoculation. C-J) 129S animals were inoculated with d42m1 tumors, and treated with rat IgG, α-PD-1 or α-PD-L1 at day 10, 13 and 16. Representative flow cytometry dot plot of mAlg8 tetramer positive cells isolated from the tumor of C) α-PD-1 treated (clone G4) or D) α-PD-L1 (clone 10F.9G2) treated animals the day after the 3rd treatment. Total tumor-infiltrating E) CD8 T cells, F) mAlg8+ tetramer + G) mLama4+ tetramer + T cells isolated from α-PD-1 treated animals. In independent experiments, d42m1-T3 tumor bearing animals were treated on the same treatment schedule as in (C-G) with α-PD-L1 and total H) CD8, I) mAlg8+, and J) mLama4+ T cells were quantified. K-M) MCA-205 tumor bearing Nur77-GFP were treated starting on day 10 as in C-J. K) Representative flow plots showing PD-1 and Nur77 expression in the tumor and spleen of MCA-205 tumor bearing Nur77-GFP mice treated with rat IgG the day after the 3rd treatment. L) Quantification of percent Nur77-GFP hi CD8 T cells in the tumors and M) spleen after the 3rd treatment. A-B) Data is representative of more than 6 independent experiments with more than 3 animals per experiment. C,E-G) Data is representative of 3 independent experiments with 3 animals per treatment/experiment. B,H-J) Representative data of at least 3 independent experiments with 3-5 animals per treatment/experiment. K-N) Data is representative of two or three independent experiments with 2-5 animals/treatment/experiment.

We then evaluated the frequency and total number of T cells in tumor-bearing animals treated with either anti-PD-1 (clone G4) or anti-PD-L1 (clone 10F.9G2). The total frequency of tumor-antigen specific CD8 T cells in the tumor was not increased after PD-1 blockade (Fig 2C) but was increased after anti-PD-L1 treatment (Fig 2D). Moreover, after anti-PD-1 treatment, there was a dramatic decrease in total CD8 T cells (Fig 2E) as well as tetramer+ T cells (Fig 2F-G). This observation was specific to the tumor microenvironment as there was an increase in the total number of CD8 T cells and tetramer positive T cells in the tumor draining lymph node after treatment with anti-PD-1 (Supplemental Fig 2). In contrast, anti-PD-L1 treatment lead to an increase in total CD8 T cells as well as tumor specific T cells in the tumor (Fig 2H-J). We repeated these experiments with a tumor expressing the model antigen ovalbumin (MCA-OVA) and used SIINFEKL tetramers to identify endogenous tumor-antigen specific T cells. Again, in animals treated with anti-PD-1, we observed a reduction in total CD8+ T cells and tetramer+ T cells in the tumor but not the draining lymph node (Supplemental Fig 3).

Using an independent mouse model, in which tumor-specific T cells are identified by high expression of the TCR-induced Nur77GFP signal (Moran et al., 2011; Moran et al., 2016; Polesso et al., 2019) and co-express high PD-1 in the tumor but not the spleen (Fig 2K), we observed an overall decrease in the frequency of CD8+Nur77GFP^hi^ T cells in the tumor after anti-PD-1 (Fig 2L). This reduction occurred in the tumor, but not the spleen (Fig 2M), and there was no reduction when using anti-PD-L1. Taken together, we observe a decrease in the number of tumor-infiltrating T cells, in particular CD8+ tumor-antigen specific T cells, after treatment with anti-PD-1 (clone G4) but not anti-PD-L1 (clone 10F.9G2). This loss of antigen specific T cells is restricted to the tumor microenvironment and not observed in the draining lymph node or spleen of tumor-bearing animals.

### Loss of antigen-specific T cells is more pronounced with PD-1 clone G4 than RMP1-14

It is well accepted that T cells can upregulate PD-1 upon TCR ligation in inflammatory settings, including viral infections. Cytomegaloviruses (CMV) are ubiquitous herpesvirus family members that infect most mammalian species and induce robust CD8 T cell responses (Munks et al., 2006; Snyder et al., 2011; Sylwester et al., 2005). Using murine CMV (MCMV)-infected mice, we treated with anti-PD-1 early after infection (Fig 3A) to determine if we could enhance the total size of the antigen-specific CD8 T cell response. We observed that a large fraction of antigen experienced CD8 T cells (CD8+CD44^hi^) expressed PD-1 (Fig 3B). Using tetramers to identify the virus-specific CD8 T cell population (M38), we observed high expression of PD-1 in virus-specific T cells (Fig 3C). When we compared the total number of antigen experienced CD8+CD44^hi^ T cells after infection and treatment with anti-PD-1 or anti-PD-L1, we observed similar numbers after rat IgG or anti-PD-1 treatment, but a significant increase after anti-PD-L1 (Fig 3D). The total number of splenocytes was only increased after anti-PD-L1 (Supplemental Fig 4A), suggesting that the decrease in CD8+CD44^hi^ T cells was not due to a global loss of cells in these animals. Moreover, bulk splenocytes were stimulated with MCMV specific peptides (see Fig 3A for timing) and IFNγ measured. Within the total splenocyte population, there was not a significant decrease in IFNγ+ CD8 T cells (supplemental Fig 4B). However, there was a significant decrease in IFNγ+M38-specific CD8 T cells after treatment with the PD-1 clone G4 but not with clone RMP1-14 or PD-L1 antibody (Fig 3E-F). Importantly, there was a wide spread in the data from RPM1-14-treated mice in multiple independent experiments. This observation was also true for other MCMV-specific T cell epitopes (Supplemental Fig 4C-F). We also observed a significant decrease in the total number of M38-specific (Fig 3G) CD8 T cells in the spleen of clone G4 treated but not anti-PD-L1 treated animals. Taken together, these data highlight that anti-PD-L1 and anti-PD-1 inhibition differentially impact the abundance of PD-1+ virus-specific CD8 T cells.

**Figure 3.**
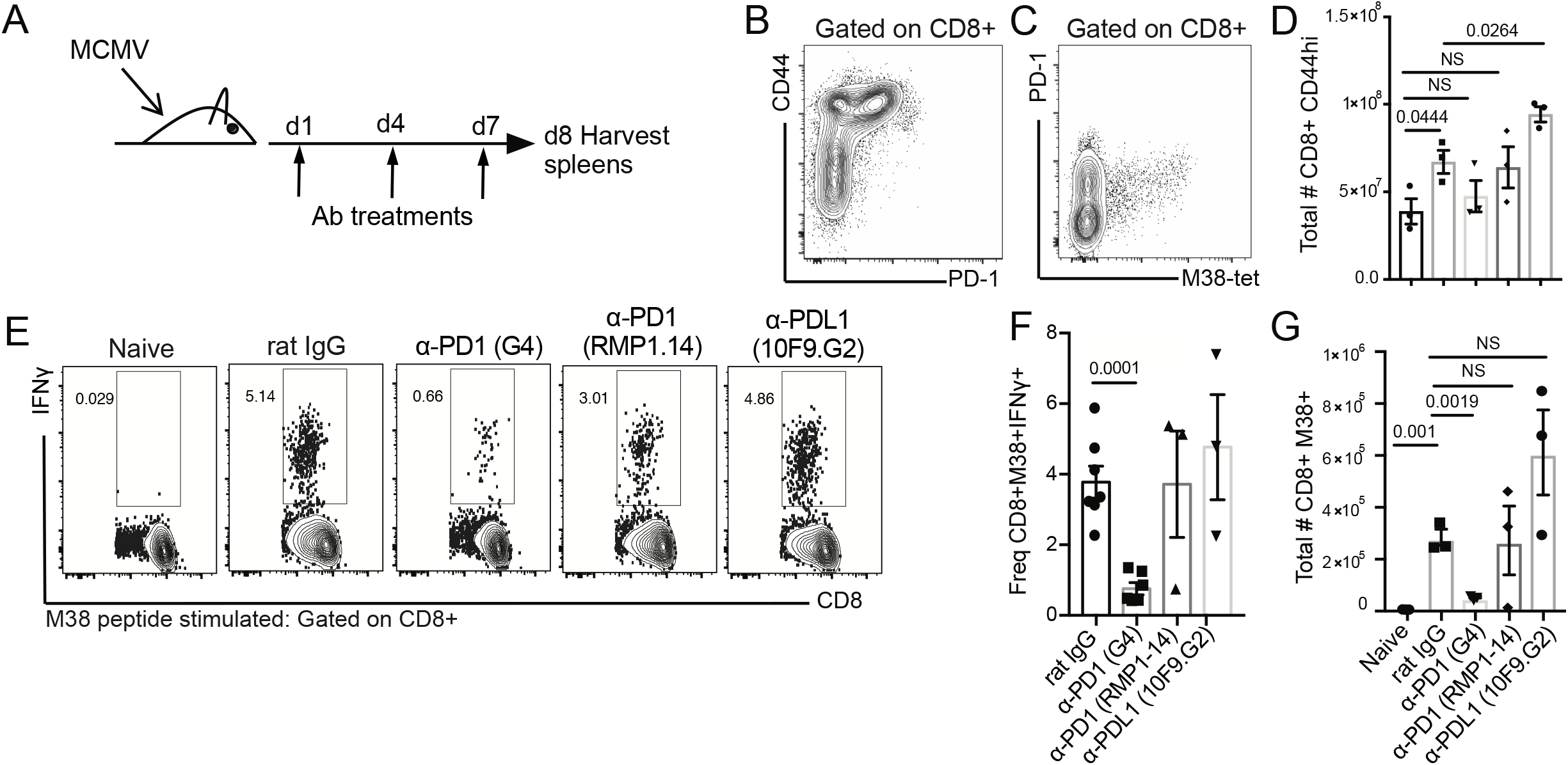
Loss of MCMV-specific CD8 T cell responses after treatment of infected animals with anti-PD-1 clone G4. A) Schematic for MCMV infection and treatment with PD-1 or PD-L1 pathway blockade. Representative cytogram of B) CD8 T cells and C) CD8 M38 tetramer + T cells from d8 spleen of rat IgG control treated animals. D) Total number of antigen experienced CD8 T cells from each treatment group. E) IFNγ intracellular cytokine staining after M38 peptide stimulation. F) Summary of the frequency of IFNγ+CD8+M38+ T cells. G) Total number of MCMV-M38 specific CD8 T cells in the spleen of each treatment group. B-G) Data is representative of two experiments with 3 animals per treatment group.

### Cross-Competition Between Anti-PD-1 Clones

PD-1 is expressed as an ‘activation marker’ on CD8 T cells after antigen recognition, as well as by exhausted antigen specific cells. Hence, in the tumor microenvironment, PD-1 expression is often used as a surrogate marker for tumor-specific T cells in the absence of known neo-antigens and tetramer tools. As noted above, MCMV infection resulted in PD-1 expression on CD8 T cells that were either CD44^hi^, or M38 tetramer+ (Fig 3A and B). However, when MCMV-infected mice were treated with anti-PD-1 (clone G4), CD8 T cells that were either CD44^hi^ or M38 tetramer+ no longer had detectable PD-1 when stained with clone 29F.1A12 (data not shown). This suggested to us that therapeutic antibody binding to PD-1 prevented its subsequent detection by a second antibody, even though the detection antibody was a different clone. There are many anti-PD-1 mAb clones in use, for both PD-1 blockade and PD-1 detection, but there are only fragmented analyses of which clones directly compete with each other. This makes it challenging to quantify the expansion and/or deletion of tumor-associated antigen specific T cells. As a resource, we tested 5 anti-PD-1 therapeutic clones in a FACS-based *in vitro* blocking assay, with the goal of identifying at least one combination of clones where the therapeutic blocking clone did not prevent FACS detection with the staining clone.

EL4 cells constitutively express high levels of PD-1, thus we stained EL4 cells with anti-PD-1 clones 29F.1A12, J43, G4, RMP1-14 and RMP1-30. Each clone detected PD-1, as expected, although the staining intensity varied widely between clones (Figure 4, top row). 29F.1A12 staining was brightest, RMP1-14 staining was weakest, while J43, G4 and RMP1-30 each stained with intermediate intensity. In the same experiment, we performed cross-blocking with unconjugated versions of the same clones (Fig 4 bottom rows). 29F.1A12 is reported to prevent PD-1 from interacting with PD-L1, and these data show that it also completely prevents PD-1 detection with nearly all other clones (Fig 4, second row). J43 and G4 also competed with most other clones. In contrast, RMP1-14 also blocks PD-L1 (Totsuka et al., 2005; Yamazaki et al., 2005), but did not interfere with detection of PD-1 by any other clone (Fig 4, 5th row). These data, summarized in supplemental Table I, suggest that RMP1-14 could be used for immunotherapy *in vivo* without preventing detection with any other clone, with consideration of the data spread presented (Fig 3).

**Figure 4.**
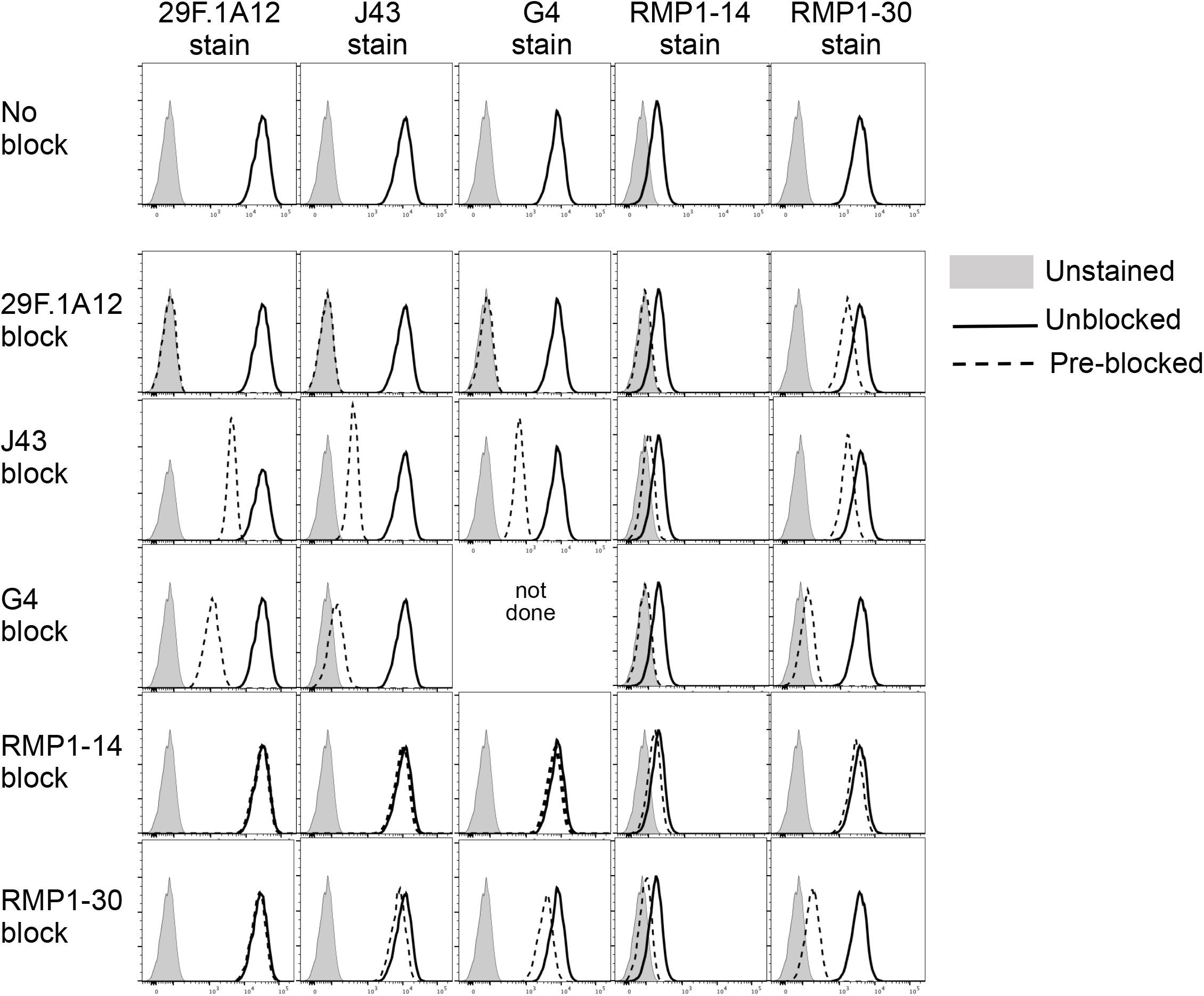
Cross-Competition Between Anti-PD-1 Clones. Top Row) PD-1 was detected on EL4 cells with αPD-1 clones, at 1 ug/ml, as indicated above. Bottom Rows) In the same experiment, some EL4 cells were pre-incubated with 10 ug/ml of blocking mAb, as indicated on the left. (nd) indicated not done because the secondary reagent would detect both the blocking mAb and the staining mAb. Data representative of two experiments.

## Discussion

In both humans and mice, T cells activated in the tumor microenvironment frequently express PD-1, which acts to restrain CD8 T cell function. Thus, anti-PD-1 antibodies are widely used clinically, as well as and in basic research. We have demonstrated here, in two mouse tumor models and a viral infection model, that one such PD-1 clone (G4) depletes antigen-specific CD8 T cells reliably, while another clone (RMP1-14) may do so in some mice. We propose that these observations should be considered when evaluating the efficacy of PD-1 single and combination immunotherapy in preclinical mouse models.

Although there has been great interest and effort to develop mouse models that appropriately recapitulate the most salient features of human cancers, there has been little effort to determine whether widely-used anti-mouse PD-1 mAb clones act in the same manner as clinical therapeutics. This is somewhat surprising, given that the most-used anti-mouse PD-1 clones have isotypes that are known to confer depleting activity to other mouse reagents. For example, anti-PD-1 clones G4 and J43 are hamster IgG, the same species and isotype as anti-CTLA-4 clones 9H10 and 4F10, which are known to deplete Treg TILs *in vivo* (Simpson et al., 2013). Likewise, RMP1-14 and 29F.1A12 are rat IgG2a, the same species and isotype as clone 53-6.7 (CD8α), used to deplete CD8 T cells (Goldschmidt et al., 1988), and RMA4-5, used to deplete CD4 T cells (Kruisbeek, 1992). Finally, RMP1-30 is rat IgG2b, the same species and isotype as YTS169.4 (CD8α), used to deplete CD8 T cells (Cobbold et al., 1985) and GK1.5, used to deplete CD4 T cells (Waldor et al., 1985) *in vivo.*

Our data are consistent with prior work showing CD8 TIL depletion when rat anti-mouse PD-1 clone 4H2 (Dahan et al., 2015); or an unspecified clone (Pai et al., 2019) is expressed recombinantly with a mouse IgG2a backbone, which has high activating-to-inhibitory FcγR binding activity. These results, along with our own, caused us to reconsider the interpretation of other studies. For instance, Gordon and colleagues demonstrated that a majority of TAMs are PD-1^hi^, that the PD-1+ TAMs are enriched for markers of M2, pro-tumorigenic macrophages, and that high PD-1 engagement on TAMs limits their phagocytic capacity and anti-tumor function (Gordon et al., 2017). Not mutually exclusive to the authors’ interpretation, it is plausible that anti-PD-1 depletes some TAMs in that setting; a mechanism that could lead to an increase in CD8 T cell function within the tumor microenvironment and interpreted as ‘reinvigorating’ exhausted CD8 T cells.

CD8 TILs are not depleted *in vivo* when anti-PD-1 is expressed as recombinant mouse IgG1 (Dahan et al., 2015), which has naturally low activating-to-inhibitory FcγR binding activity. Likewise, CD8 TILs are not depleted when anti-PD-1 is expressed as recombinant mouse IgG2a engineered to lack FcγR binding (Pai et al., 2019). In preclinical mouse models of tumor immunotherapy, tumor regression is often the readout for therapy efficacy, sometimes complimented with correlative studies on the function of tumor-associated CD8 T cells. Distinguishing between tumor-infiltrating T cell intrinsic and extrinsic mechanisms of immunotherapy response and resistance are critical for improving translation of mouse studies and patient outcomes with these oncologic therapies.

We propose several changes to improve interpretation of pre-clinical PD-1 studies. First, common anti-PD-1 mAb clones should be more thoroughly characterized, side-by-side. Second, PD-1 mAbs lacking FcγR-mediated effector function could be developed, for example by expression as mouse IgG1 antibodies. Third, studies utilizing anti-PD-1 in mice should include quantitative analysis of PD-1+ cells, and thoughtful acknowledgment of alternative interpretation.

## Materials and Methods

### Animals, Tumor Models and Antibodies

C57BL/6 and ovalbumin tolerant RIP-mOVA mice were purchased from the Jackson Laboratory. 129S6 were purchased from Taconic. Nur77-GFP mice were provided by K.A. Hogquist (University of Minnesota, Minneapolis, MN), maintained as heterozygous transgene carriers. All animals were bred and maintained under specific pathogen-free conditions in the Oregon Health and Science University (Portland, OR) animal facility. Males and females were used for the 3’-Methylcholanthrene (MCA) 205 and MCA-205-OVA sarcoma studies presented. Males were used for the d42m1-T3 studies presented. All animal experimentation was approved by and performed according to the guidelines from the Institutional Animals Care and Use Committee at OHSU. Murine MCA-205 sarcoma cells (gift of Suyu Shu), MCA-205-OVA (gift of Andrew Weinberg) and d42m1-T3 (gift of Robert Schreiber) were propagated in vitro using Complete Media, RPMI 1640 (Lonza) containing 0.292 ng/ml glutamine, 100 U/ml streptomycin, and penicillin, 1X non-essential amino acids, 1 mmol/l sodium pyruvate, and 10 mmol/l HEPES (Sigma-Aldrich). All cell lines were tested and confirmed to be Mycoplasma and endotoxin-free using the MycoAlert Detection kit (Lonza) and the Endosafe-PTS system (Charles River Laboratories). EL4 cells (ATCC) were cultured in RPMI-1640 supplemented with 10% fetal bovine serum and antibiotics. All culture media reagents were purchased from Hyclone Laboratories unless noted otherwise. Control rat IgG was purchased from Sigma-Aldrich. Anti-PD-1 (G4) mAb was obtained from Leiping Chen. Anti-PD-1 (RMP1-14) and anti-PD-L1 (10F.9G2) was purchased from BioXcell. Animals were randomly assigned to treatment cohorts, and tumors were ~50 to 70 mm^2^ (by two-dimension caliper measurement) at the start of treatment. Any animal with a tumor > 200 mm^2^ was euthanized per our guidelines from the Institutional Animal Care and Use Committee. No outliers were excluded from the data presented.

### Tumor Challenge and Lymphocyte Isolation

0.5×10^6^ MCA-205 tumor cells were injected on both hind flanks of C57BL6 or Nur77-GFP mice, and 0.5×10^6^ MCA-205-OVA or 1×10^6^ d43m1-T3 tumor cells were injected on both hind flanks of RIP-mOVA or 129S6 mice respectively (minimum of 5 animals per group per survival experiments and minimum of 3 per group per phenotyping experiments). When the tumors reached ~70 mm^2^ (except for Fig 1 where animals were treated at day 5), the tumor-bearing mice were treated with 3 (phenotyping) or 4 (survival) doses of 200 ug rat IgG, anti-PD-1 or anti-PD-L1 3 days apart. All antibody injections were given i.p. For survival experiments, tumors were measured 3 times per week until the tumor reached 150 mm^2^ and animals were euthanized. For phenotyping experiments, TIL were harvested by dissection of tumor tissue into small fragments in a 50 ml conical tube followed by digestion at room temperature in a bacterial shaker at 180 rpm for 30 minutes in 1 mg/ml collagenase type IV (Worthington Biochemicals), 100 ug/ml hyaluronidase (Sigma-Aldrich) and 20 mg/ml DNase (Roche) in PBS. Cell were then further disrupted with a 1-cc syringe plunger through a 70-um nylon cell strainer (BD Biosciences), and filtered to obtain a single cell suspension. Tumor dLN (inguinal) and spleens were harvested and processed to obtain single-cell suspensions using frosted ends of microscope slides. Spleens were incubated with ammonium chloride potassium lysing buffer (Lonza) for 3 min at room temperature to lyse red blood cells. Cells were rinsed with PBS containing 1% FBS and 4 mM EDTA.

### MCMV Infection

C57BL/6 mice were infected intraperitoneally (i.p) with 1-2 x 10^5^ PFU of wild type MCMV, strain MW97.01, which is derived from a bacterial artificial chromosome of the Smith strain (Wagner et al., 1999). Virus stocks were grown and titered on BALB 3T3 cells. On day 1 and 4 post infection, mice were treated with 200 ug of rat IgG, anti-PD-1 (G4), anti-PD-1 (RMP1-14), or anti-PD-L1 (10F.9G2) i.p. On day 7, spleens were harvested, splenocytes isolated as described above and stained for flow cytometry.

### Flow Cytometry

For flow cytometry analysis, cells were incubated for 20 min on ice with e506 fixable viability dye and Fc block. Cells isolated from MCA-205-OVA tumor bearing animals were stained with H-2K^b^:OVAp (SIINFELK) MHC class I tetramer to evaluate ovalbumin specific responses, cells from d42m1-T3 tumor bearing experiments were stained with H2K^b^:mLama4 (VGFNFRTL) and H2K^b^:mAlg8 (ITYTWTRL) tetramers to evaluate tumor specific responses, and MCMV infected animals were stained with H2K^b^:M38, H2K^b^:M45 tetramers to evaluate MCMV specific responses. All MHC class I tetramers were generated by the National Institute of Health Tetramer Core Facility. Tetramer staining was performed on ice for 30 minutes in presence of Fc block. Cells were then incubated for 20 min on ice with TCRß (H57-597), CD8 (53-6.7), CD44 (IM7), PD-1 (J43) antibodies. All antibodies and viability dies were purchased from eBioscience, Biolegend, or BD Biosciences. Unless noted otherwise in the figure legend, cells were gated through live/TCRß+ gates for analysis. Cells were analyzed with an LSR II or Fortessa flow cytometer (BD Biosciences) using FlowJo software (FlowJo, LLC).

### In Vitro Activation and Intracellular Cytokine Staining

To quantitate MCMV-specific CD8 T cell responses, splenocytes were stimulated with 1 ug/ml of M38 (316-323; SSPPMFRV) synthetic peptide (Genemed Synthesis Inc.) for 5 hours in the presence of BFA. Cells were then stained for surface markers as indicated, fixed and permeabilized with the Cytofix/Cytoperm Kit (BD Biosciences), and stained for intracellular IFNγ using APC-XMG1.2 (eBioscience).

### In Vitro Blocking Assay

EL4 cells were plated in 96 well plates with 10^5^ cells per well. Blocking antibodies were added at 10 ug/ml for 30 minutes at 4°C, in PBS containing 2% FBS and 0.1% sodium azide. Blocking was performed using unconjugated 29F.1A12 (BioXcell), unconjugated J43 (eBioscience), unconjugated G4 (obtained from Leiping Chen), unconjugated RMP1-14 (BioLegend), or unconjugated RMP1-30 (BioLegend). Following the block, 1 ug/ml staining antibody was added, without removing the blocking antibody, and incubated a further 20 minutes at 4°C, before washing three times. Staining was performed using PE-labeled 29F.1A12 (BioLegend), PE-labeled J43 (Ebioscience), PE-labeled RMP1-14 (BioLegend), PE-labeled RMP1-30 (BioLegend), or unconjugated G4 followed by PE-conjugated mouse anti-hamster IgG cocktail (clones G94-90.5 and G70-204; BD). All cells were then analyzed on an LSR II (BD Life Sciences) and analyzed with FlowJo software (FlowJo LLC).

Because FACS data is visualized on a log scale, inhibition was calculated on a log scale. First, unstained GMFI values, unblocked GMFI values and blocked GMFI values were converted to log values. Percent inhibition was then calculated as: 1 – ((blocked – unstained) / (unblocked – unstained)).

### Statistical Analysis

Statistical analyses were performed using unpaired two-tailed Student’s t-test (for comparison between two groups), or one-way ANOVA for multiple comparisons, with GraphPad Prism 6 (GraphPad Software). Error bars represent SEM unless noted otherwise in the figure legend. Statistical tests and *p* values are specified for each panel in the respective figure legends, and *p* values < 0.05 were considered significant. Biological replicates (individual animals) for each experiment are indicated in the figure legends.

### Study Approval

All animal experiments were approved by the Institutional Animal Care and Use Committee of Oregon Health and Science University in Portland, OR.

