## Supplemental Figure 1 for "PD-1 Specific ‘Blocking’ Antibodies That Deplete PD-1+ T Cells Present An Inconvenient Variable In Pre-clinical Immunotherapy Experiments"

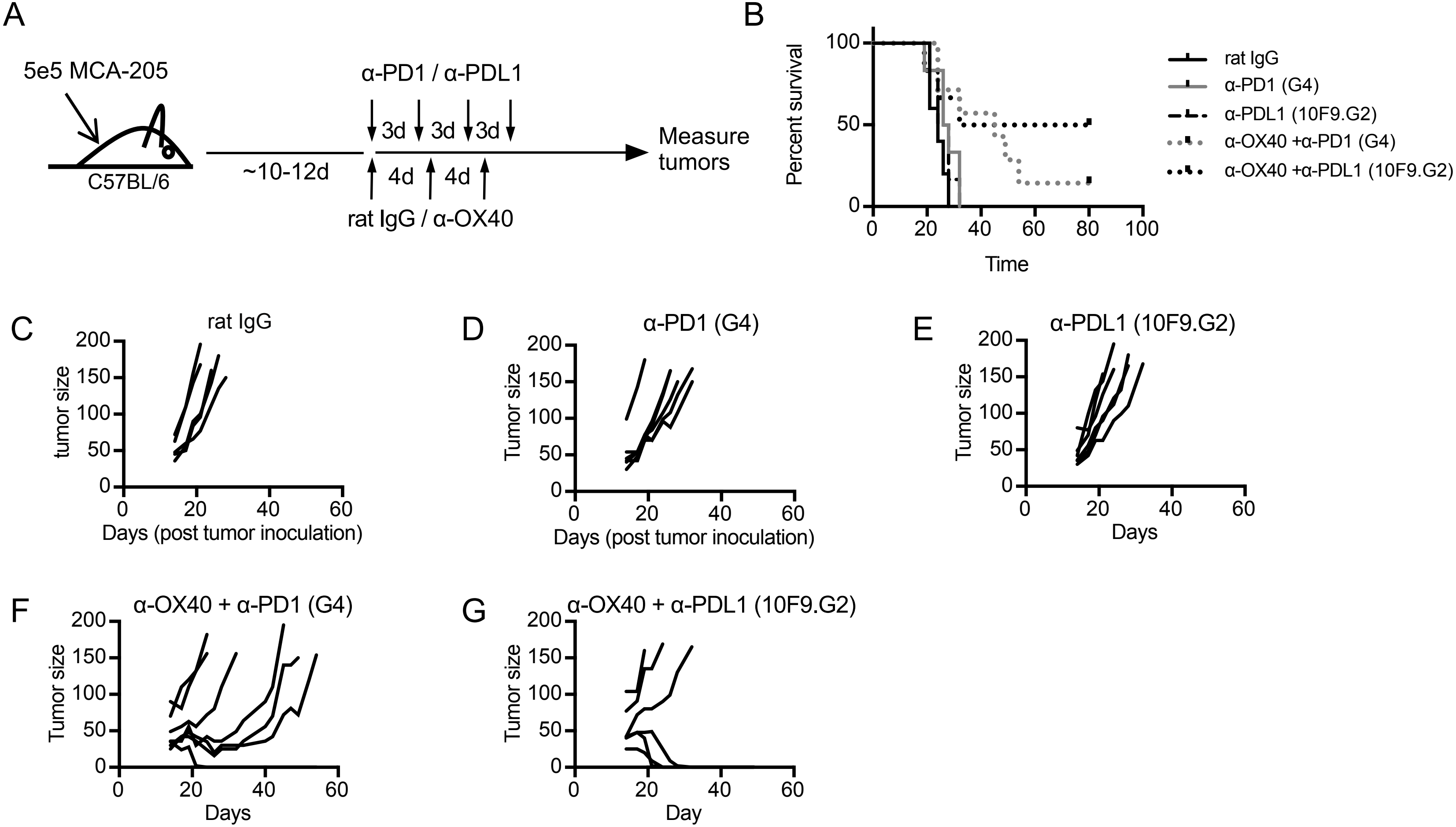


**Supplemental Figure 1. Survival outcomes are different in combination immunotherapy with anti-PD-1 vs anti-PD-L1 treatment.** A) Experimental design of tumor experiments. Tumors were measured 3 times a week and animals were euthanized when the tumors were >150mm2 or when tumors became ulcerated. B) Overall survival of MCA205 tumor-bearing C57Bl/6 mice. C-G) Individual tumor growth curves for each treatment group. Data are representative of more than 3 independent experiments with 5-7 animals per treatment group.
