## Supplemental Figure 2 for "PD-1 Specific ‘Blocking’ Antibodies That Deplete PD-1+ T Cells Present An Inconvenient Variable In Pre-clinical Immunotherapy Experiments"

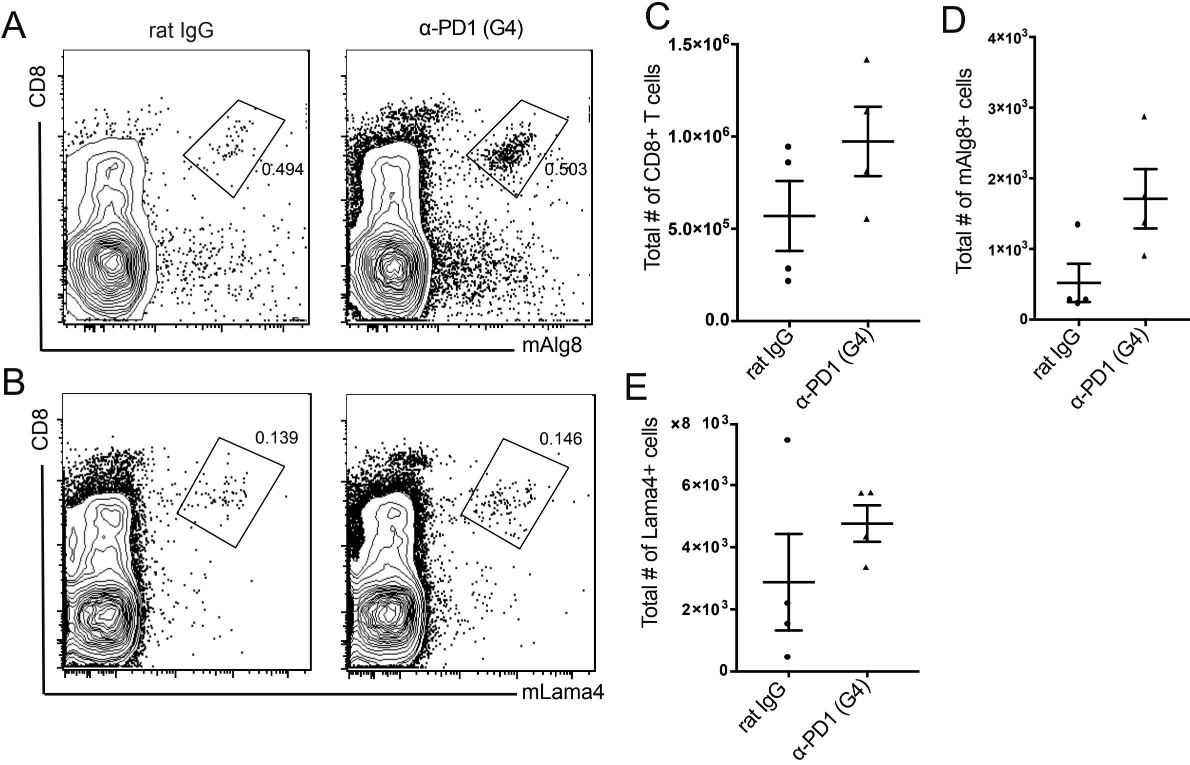


**Supplemental Figure 2. Tumor antigen specific T cells numbers are similar in the draining lymph node in rat IgG control and anti-PD1 treated animals.** Representative flow cytogram from d42m1-T3 tumor draining lymph node stained with A) mAlg8 and B) mLama4 tetramer. Summary of total numbers of C) total CD8 T cells, D) mAlg8+ and E) mLama4+ CD8 T cells in the tumor draining lymph node of rat IgG or anti-PD-1 (clone G4) treated animals.
