## Supplemental Figure 3 for "PD-1 Specific ‘Blocking’ Antibodies That Deplete PD-1+ T Cells Present An Inconvenient Variable In Pre-clinical Immunotherapy Experiments"

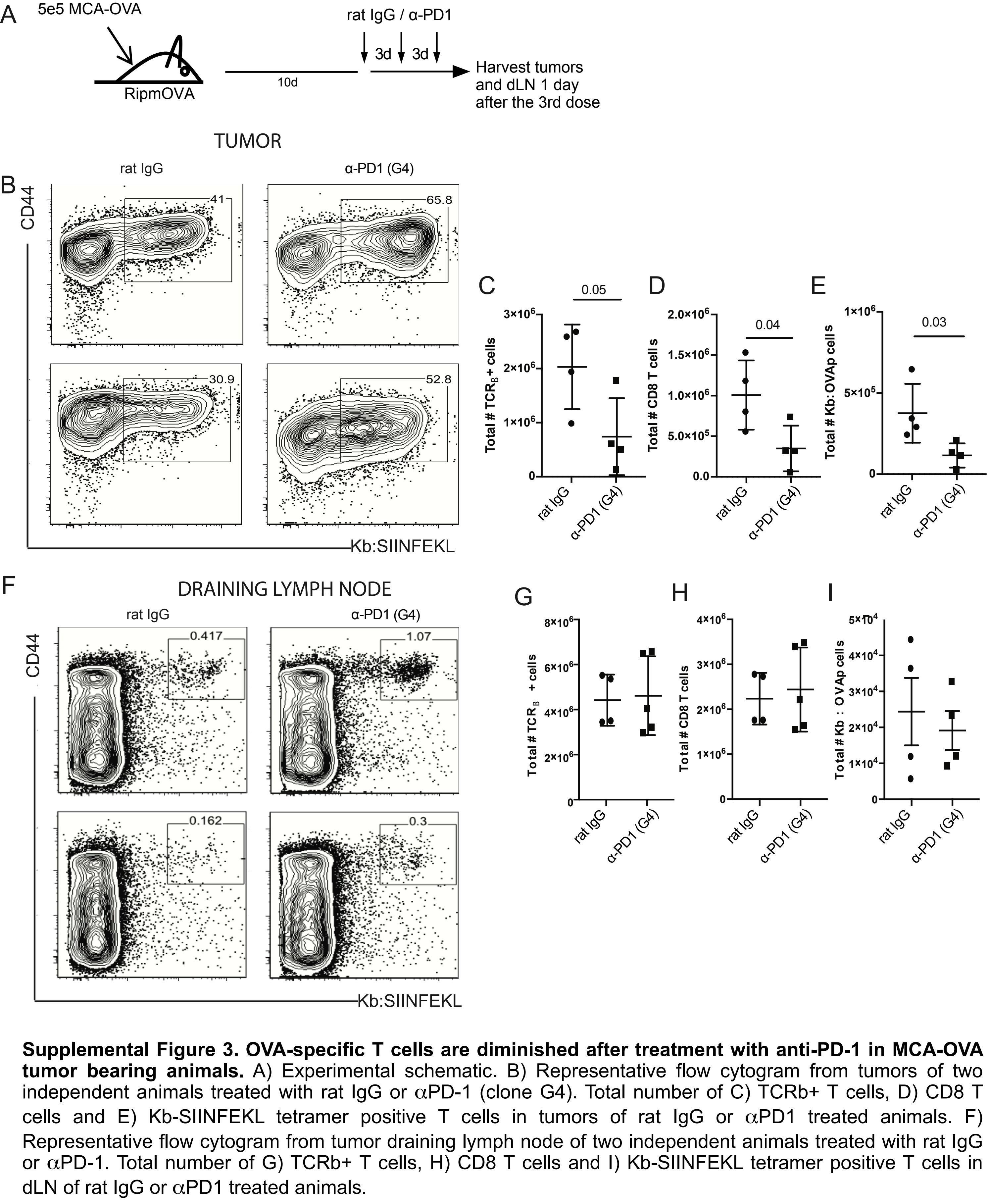


**Supplemental Figure 3. OVA-specific T cells are diminished after treatment with anti-PD-1 in MCA-OVA tumor bearing animals.** A) Experimental schematic. B) Representative flow cytogram from tumors of two independent animals treated with rat IgG or $\alpha$PD-1 (clone G4). Total number of C) TCRb+ T cells, D) CD8 T cells and E) Kb-SIINFEKL tetramer positive T cells in tumors of rat IgG or $\alpha$PD1 treated animals. F) Representative flow cytogram from tumor draining lymph node of two independent animals treated with rat IgG or $\alpha$PD-1. Total number of G) TCRb+ T cells, H) CD8 T cells and I) Kb-SIINFEKL tetramer positive T cells in dLN of rat IgG or $\alpha$PD1 treated animals.
