## Supplemental Figure 4 for "PD-1 Specific ‘Blocking’ Antibodies That Deplete PD-1+ T Cells Present An Inconvenient Variable In Pre-clinical Immunotherapy Experiments"

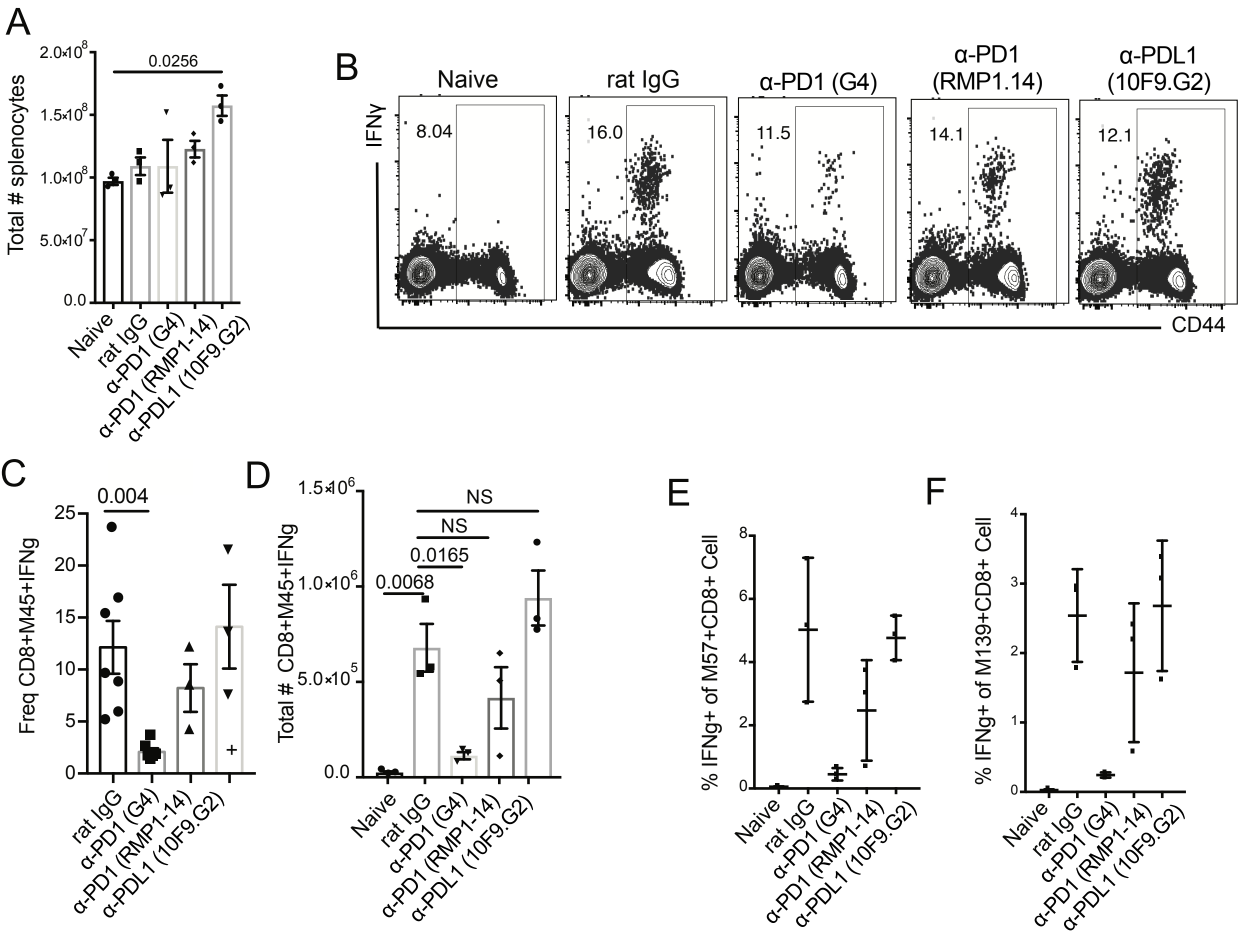


**Supplemental Figure 4. Loss of MCMV-specific CD8 T cell responses after treatment of infected animals with anti-PD-1 clone G4.** A) Total number of splenocytes between each treatment group. B) Representative flow cytogram of MC38+IFNγ+ T cells after peptide stimulation and intracellular cytokine staining. C) Frequency of M45+IFNγ+ T cells after peptide stimulation. D) Total number of M45+ T cells in the spleen after MCMV infection. Frequency of IFNγ+ E) M57 specific and F) M139 specific CD8 T cells in the spleen after peptide stimulation.
