## Supplemental Table 1 for "PD-1 Specific ‘Blocking’ Antibodies That Deplete PD-1+ T Cells Present An Inconvenient Variable In Pre-clinical Immunotherapy Experiments"

**Supplemental table 1:** Cross-competition between anti-PD-1 clones

| | 29F.1A12 (PE) | J43 (PE) | G4 (PE) | RMP 1-14 (PE) | RMP 1-30 (PE) |
|---|---|---|---|---|---|
| 29F.1A12 (block) | 99% | 100% | 98% | 90% | 20% |
| J43 (block) | 33% | 65% | 53% | 64% | 21% |
| G4 (block) | 54% | 86% | N.D. | 90% | 85% |
| RMP 1-14 (block) | 0% | 2% | 4% | 24% | 6% |
| RMP 1-30 (block) | 2% | 7% | 16% | 66% | 75% |

Numbers indicate percent inhibition of each blocking antibody (row) for each staining antibody (column).

N.D. = not done, because the detection antibody would recognize both the blocking antibody and the staining antibody.
